# The human hippocampus retroactively binds experiences across time

**DOI:** 10.64898/2026.10.20.760394

**Authors:** Xianhui He, Marit Petzka, Mohammad F. Khazali, Armin Brandt, Marcel S. Kehl, Lukas Kunz, Andreas Schulze-Bonhage, Bernhard P. Staresina

## Abstract

Creating memories requires linking events across seconds and minutes, but canonical mechanisms of associative plasticity operate over much shorter timescales. This poses a fundamental problem: how can the brain bind information that arrives at different times? Computational model and animal studies implicate the hippocampus in associating events across time, but when the hippocampus becomes engaged and how it interacts with representations of preceding events are unknown. Here we show, using intracranial recordings in humans forming image–word associations across a ∼2-s temporal gap, that hippocampal high-frequency activity predicts later memory when the second event arrives, but not during the initial image or intervening delay. At this moment, neurons tuned to the preceding image reinstate stimulus-specific firing despite its sensory absence. Across trials, hippocampal high-frequency activity was selectively coupled to this reinstatement during successful encoding and increased before spiking of image-tuned neurons. These findings reveal a retroactive mode of hippocampal binding, whereby representations of preceding events are recruited when new information arrives. This mechanism provides a neural solution for binding experiences across the temporal discontinuities that characterize everyday life.

## Introduction

You and a fellow birdwatcher glimpse a bird before it disappears into the trees. Moments later, your companion identifies it as a chiffchaff. Learning the bird’s identity requires the brain to associate information arriving now (“chiffchaff”) with a representation of an event that has already passed (the visual representation of the bird). Episodic memory routinely joins elements that were never simultaneously present in the environment (Tulving, 1983, 2002). However, canonical Hebbian plasticity mechanisms, including spike-timing-dependent plasticity, depend on pre- and postsynaptic activity occurring within tens of milliseconds (Dan & Poo, 2004; Feldman, 2012), far shorter than the temporal gaps that often separate elements of an experience. How the brain brings such temporally discontiguous information into a common window for associative binding thus remains a major puzzle in memory research (Clewett et al., 2019; Davachi, 2006; Eichenbaum, 2004).

The hippocampus is a prime candidate for solving this problem. Lesion studies across species have shown that hippocampal integrity is particularly important for trace conditioning, in which the conditioned and unconditioned stimuli are separated in time, whereas conventional delay conditioning, in which the stimuli overlap, can proceed without an intact hippocampus (Clark & Squire, 1998; Moyer et al., 1990; Solomon et al., 1986). Computational models likewise suggest that the hippocampal architecture is well suited to bridge discontinuities between event elements (Lisman, 1999; Treves & Rolls, 1994; Wallenstein et al., 1998; Whittington et al., 2020). In humans, fMRI studies have implicated the hippocampus and adjacent medial temporal lobe regions in forming associations across temporal gaps (Hales & Brewer, 2011; Hales et al., 2009; Qin et al., 2007; Staresina & Davachi, 2009), but the temporal ambiguity of the BOLD signal leaves the underlying mechanism unresolved. Various scenarios are conceivable. First, hippocampal engagement might already distinguish successful vs. unsuccessful encoding when the first stimulus is encountered. Second, the hippocampus might maintain the preceding event during the intervening delay, acting as a mnemonic buffer. Third, hippocampal involvement might emerge only when the second stimulus arrives, retroactively bringing the current and preceding event representations together for binding. Human intracranial recordings provide the temporal resolution needed to adjudicate between these three scenarios, with previous work showing that hippocampal high-frequency activity (HFA; ∼50–150 Hz) tracks successful memory formation and retrieval on a subsecond timescale (Burke et al., 2014; Sederberg et al., 2007; Staresina et al., 2016; Staresina & Wimber, 2019; Treder et al., 2021).

Pinpointing when the hippocampus engages, however, addresses only half of the binding problem: the hippocampus must also interact with representations of the information to be bound. Human single-neuron recordings provide a cellular readout of such content, having revealed neurons that respond selectively and invariantly to particular stimuli or concepts and can reinstate these representations in their sensory absence (Gelbard-Sagiv et al., 2008; Kehl et al., 2026; Mackay et al., 2024; Quiroga et al., 2005; Staresina et al., 2019). Whether hippocampal activity couples to such stimulus-specific firing at the moment of binding, and whether this interaction predicts subsequent memory, is unknown.

To determine when hippocampal binding occurs and how it engages with the content of an unfolding episode, we combined macroelectrode with neuronal spiking recordings from the human medial temporal lobe while participants associated an image with a verb across an intervening ∼2-s delay (Fig. 1a). We show (i) that hippocampal HFA predicts memory success specifically when the second event arrives, but not during initial image presentation or the intervening delay; (ii) that image-tuned neurons reinstate the preceding stimulus at this same moment; and (iii) that hippocampal HFA couples to this reinstated firing selectively during successful encoding, rising before spikes in image-tuned units. Together, these findings reveal a retroactive hippocampal binding process that recruits representations of preceding events when new information arrives, thereby bridging temporal gaps during associative memory formation.

**Figure 1.**
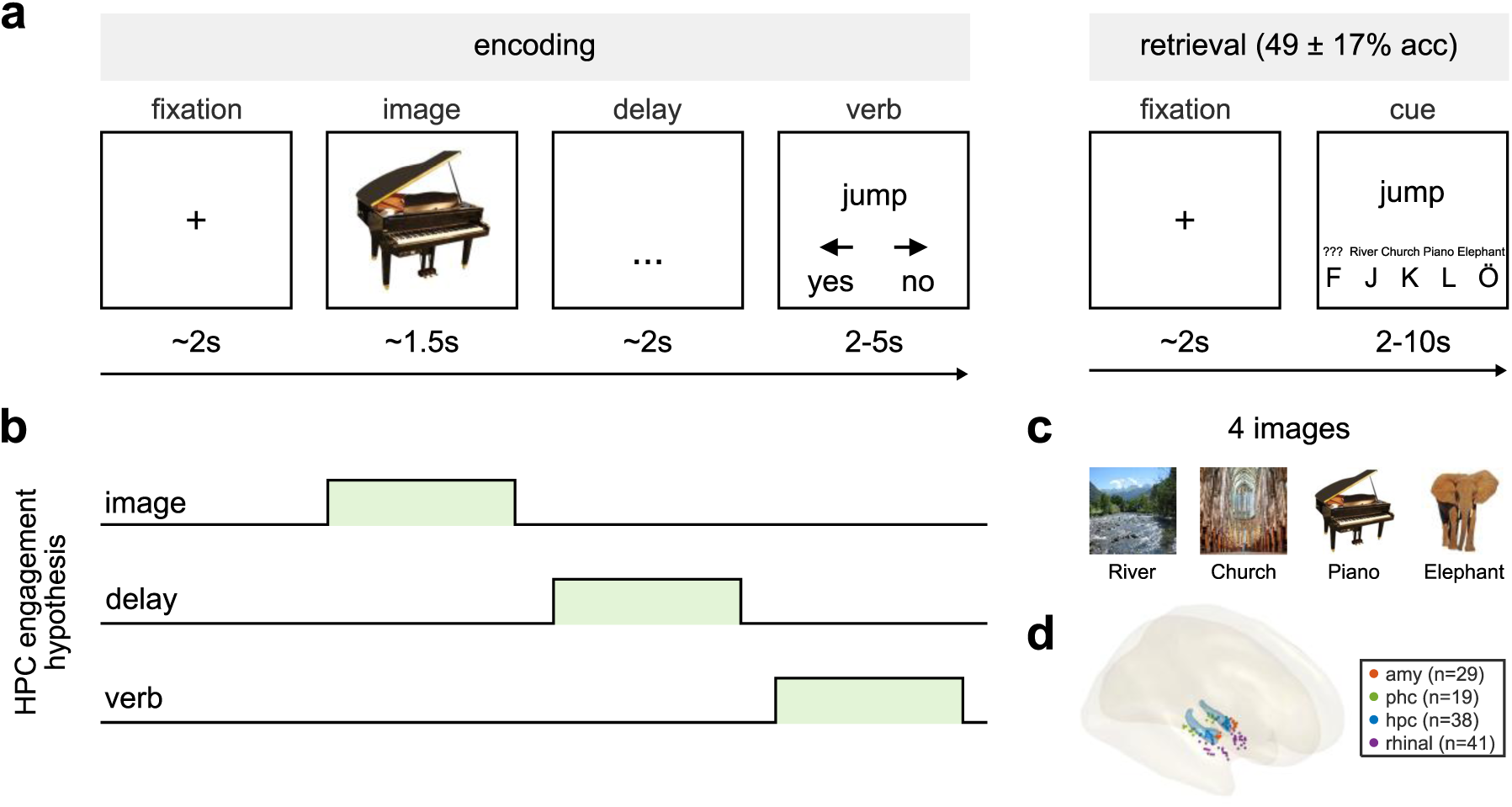
Task design and intracranial coverage of the medial temporal lobe. **a)** Trial structure. During encoding (114 ± 13 trials per participant, mean ± s.d.), each trial consisted of a fixation cross (∼2 s), one of four images (1.5–1.8 s), a blank delay (1.75–2.50 s) and a verb (2–5 s; minimum 2 s on screen irrespective of the response). Participants judged whether the image–verb association was plausible (left, yes/plausible; right, no/implausible). During retrieval (49 ± 17% accuracy, mean ± s.d., 25% chance level), a fixation cross (∼2 s) was followed by the verb cue (2–10 s; minimum 2 s on screen irrespective of the response), after which participants selected the image previously paired with the verb (images indicated by letters and German words; F, forgotten/no response). **b)** Candidate temporal profiles of hippocampal engagement during encoding: selectively recruited at image onset, during the delay or at verb onset. **c)** The four images used during encoding (river, church, piano and elephant), each mapped to a response letter. **d)** Pooled intracranial EEG (iEEG) contact locations across participants (N = 15) in the amygdala (Amy), parahippocampal cortex (PHC), hippocampus (HPC) and entorhinal/perirhinal cortex (rhinal cortex). The hippocampal surface is shown in blue.

## Results

Fifteen patients undergoing intracranial EEG monitoring performed an image-verb associative memory task in which the to-be-associated elements (image and verb) were separated by a temporal gap (Fig. 1a, b). On each encoding trial, one of four images (river, church, piano, elephant; Fig. 1c) was presented for 1.5-1.8 s, followed by a blank delay of 1.75-2.50 s and then by a verb. Participants judged whether the image-verb pairing was plausible (114 ± 13 trials per participant; mean plausibility reaction time (RT) 2.5 ± 0.5 s). Because the verb appeared only after the image had been removed from the screen and after an intervening delay, the earliest moment at which the two elements could be bound was temporally separated from both image presentation and delay maintenance. The three scenarios outlined above therefore predicted subsequent-memory effects at three different moments within the trial — at image onset, across the delay, or at verb onset (Fig. 1b). In a subsequent retrieval period, memory for each pairing was then probed with the verb as a cue and the four images as response options along with a ‘don’t know’ option (49 ± 17% correct; chance level, 25%; one-sample *t*-test against chance: *t*(14) = 5.50, *p* < 0.001). We recorded 127 MTL contacts across all 15 participants (hippocampus, n = 38; amygdala, n = 29; parahippocampal cortex, n = 19; rhinal cortex, n = 41; Fig. 1d), together with 440 MTL units in the 12 participants who additionally had microwires implanted (Table S1).

### Hippocampal HFA indicates successful encoding after verb onset

Our data unambiguously favoured one of these predictions. Specifically, hippocampal high-frequency activity (HFA, 50-150 Hz) separated later-remembered (R) from later-forgotten (F) trials only after verb onset. A time-resolved contrast across hippocampal contacts (n = 38) revealed significant later-remembered > later-forgotten clusters from around 0.25 s to 3 s after verb onset (*p*s = 0.006, contact-level linear mixed-effects (LME) cluster-based permutation test, Bonferroni corrected across the three periods, i.e., image, delay and verb, see Methods). No significant differences were observed during the image or the delay period (*p*s > 0.642, Bonferroni corrected; Fig. 2a). The same pattern was obtained without restricting the analysis to a predefined frequency band: in the full time-frequency decomposition (2-200 Hz), the only significant subsequent-memory cluster was confined to the verb period, spanning 25 to 180 Hz with its peak in the high-frequency gamma range (*p* = 0.006, Bonferroni corrected; Fig. S1). Comparing period-averaged HFA directly confirmed this dissociation. Verb-period HFA (from verb onset to trial-specific RT) was higher on R than on F trials, whereas the image and delay periods showed no such separation, yielding significant *period* (image/delay, verb) × *memory* (R, F) interactions for verb versus image (*p* < 0.001) and for verb versus delay (*p* < 0.001; n = 38 contacts; Fig. 2b). This difference did not reflect a longer time-on-task window during verb processing, as plausibility RTs did not differ between R and F trials (*t*(14) = 0.31, *p* = 0.759).

**Figure 2.**
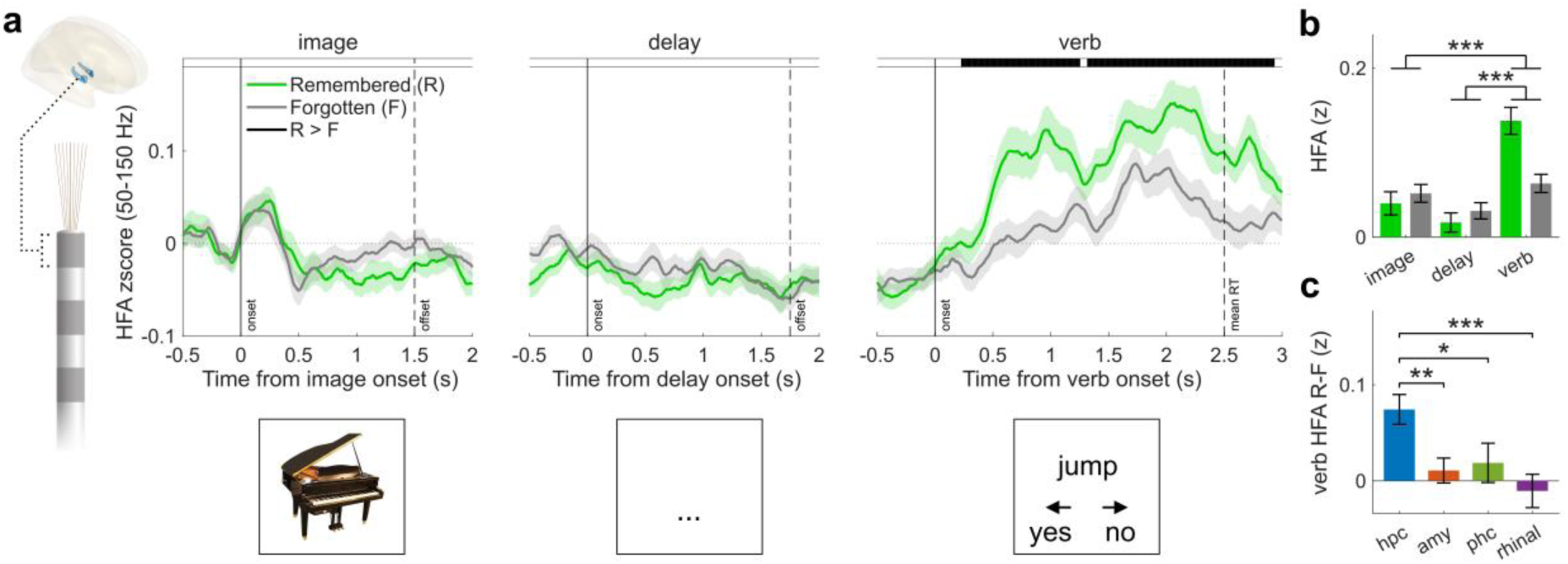
Subsequent Memory Effect in the hippocampus at verb onset. **a)** Left, schematic of a hippocampal iEEG contact. Main panels, time courses of hippocampal high-frequency activity (HFA; 50–150 Hz; z-scored across all concatenated encoding periods) for later-remembered (R) and later-forgotten (F) trials, aligned to image, delay and verb onset periods. Lines and shading indicate mean ± s.e.m. across hippocampal contacts (n = 38). Solid vertical lines indicate event onset; dashed vertical lines indicate minimum image offset (1.5 s), delay offset (1.75 s) or mean plausibility-judgement reaction time (RT; 2.5 s) for the verb period. The horizontal black bar indicates a significant R > F cluster (contact-level linear mixed-effects (LME) cluster-based permutation test: HFA ∼ memory + (1|subject) + (1|subject:contact), 1,000 permutations, *p* < 0.05, Bonferroni corrected across the three periods). No significant cluster was detected during the image or delay period. **b)** Mean HFA for R and F trials during the image ([0, 1.5] s), delay ([0, 1.75] s) and verb ([0, RT]) periods. Bars indicate mean ± s.e.m. across hippocampal contacts (n = 38). Brackets indicate period × memory interactions from 2 × 2 LME models (HFA ∼ period × memory + (1|subject) + (1|subject:contact)). \*\*\**p* < 0.001. **c)** Mean verb-period HFA difference (R - F) across MTL regions. Bars indicate mean ± s.e.m. across contacts (hippocampus, n = 38; amygdala, n = 29; parahippocampal cortex, n = 19; rhinal cortex, n = 41). The effect of region was assessed using an LME model (R-F ∼ region + (1|subject)). The bracket shows the contrast of hippocampus against each of the other three regions (Bonferroni corrected). \**p* < 0.05, ** *p* < 0.01, \*\*\**p* < 0.001.

We next probed the hippocampal specificity of this effect by examining the three other MTL regions (amygdala, n = 29; parahippocampal cortex, n = 19; rhinal cortex, n = 41). No significant difference between R and F emerged in any period in any of these regions (*p*s > 0.162; Fig. S2). Likewise, when averaging HFA across the entire verb-period (verb onset to RT), the R minus F difference was not significant in any of these MTL regions (*p*s > 0.552, Bonferroni corrected). Direct comparisons across regions confirmed that the verb-period subsequent memory effect was specific to the hippocampus: the R − F difference was larger in the hippocampus than in each of the other three MTL regions individually (*p*s < 0.034, Bonferroni corrected; Fig. 2c). In sum, these results identify a hippocampus-selective subsequent-memory effect after the entire episode had unfolded.

### Image-tuned units reinstate their preferred stimulus during the verb period

These results identify a hippocampal signal associated with successful binding in the final period of the trial, i.e., when all episodic elements (image and word) have been encountered. However, how does this binding mechanism interact with the to-be-bound content, i.e., the image entity that needs to be associated with the current verb? To recover the image-specific content available at that moment, we identified image-tuned neurons (‘units’) in a separate one-back ‘localiser’ task in which the same four images were presented without any associative/memory demand (Fig. 3a). Following established criteria (Kunz et al., 2024), a unit was classified as image-tuned only if its firing rate (FR) during its preferred image exceeded the FR during the non-preferred images in a sustained manner (Fig. 3b). This yielded 77 of 440 units (17.5%), distributed across MTL regions (rhinal cortex, n = 27; hippocampus, n = 23; parahippocampal cortex, n = 19; amygdala, n = 8) and approximately balanced across the four preferred images (17-23 units each; Fig. 3c).

**Figure 3.**
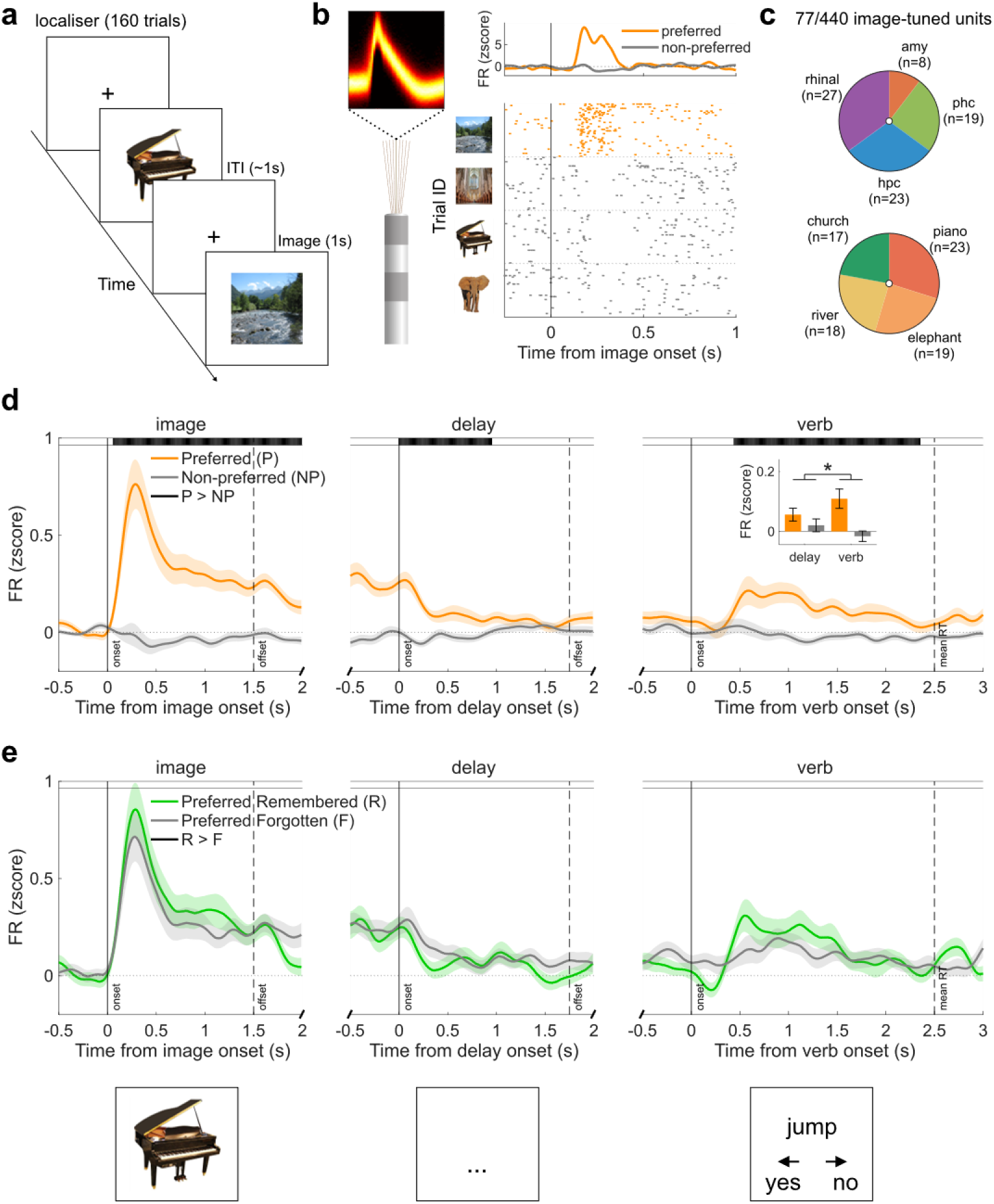
Image-tuned MTL units reactivate after verb onset. **a)** One-back ‘localiser’ task used to identify image-tuned units (160 trials; 1-s image presentation followed by ∼1-s intertrial interval). Participants responded to immediate image repetitions. **b)** Example image-tuned unit (right rhinal cortex). Top left, spike-waveform density (n = 14,931 spikes). Top right, mean baseline-z-scored firing rate for preferred (orange) and non-preferred (grey) image trials. Bottom, spike raster grouped by localiser image (river, church, piano, elephant, from top to bottom). Image tuning was defined as a sustained, significantly higher firing rate evoked by the preferred image than by non-preferred images following image onset. **c)** Distribution of image-tuned units (77 of 440 units from 11 participants; 17.5%). Top, distribution across MTL regions: amygdala (n = 8 of 156 units; 5.1%), parahippocampal cortex (n = 19 of 55 units; 34.5%), hippocampus (n = 23 of 124 units; 18.5%) and rhinal cortex (n = 27 of 105 units; 25.7%). Bottom, distribution by preferred image: piano (n = 23), elephant (n = 19), river (n = 18) and church (n = 17). **d)** Baseline-z-scored firing rates of image-tuned units on preferred (P) and non-preferred (NP) trials, aligned to image, delay and verb onset. Lines and shading indicate mean ± s.e.m. across units (n = 77). The horizontal black bar indicates a significant P > NP cluster (unit-level LME cluster-based permutation test: FR ∼ image + (1|subject) + (1|subject:region:unit), 1,000 permutations, *p* < 0.05, Bonferroni corrected across the three periods). Inset, mean firing rate during the late delay (1 s preceding verb onset) and verb ([0, RT]) periods for P and NP trials. Bars indicate mean ± s.e.m. across units (n = 77). Brackets indicate period × image interactions from 2 × 2 LME models (FR ∼ period × image + (1|subject) + (1|subject:region:unit)). \**p* < 0.05. **e)** Same as in (d) but for preferred, later-remembered (R) and preferred, later-forgotten (F) trials. No significant cluster was found (unit-level LME cluster-based permutation test, *p* > 0.05, Bonferroni corrected across the three periods).

The FR of these image-tuned units was then used to provide a readout of image-specific content during the different trial periods. As expected, FR were markedly elevated for the preferred vs. non-preferred image during the initial image presentation (*p* = 0.006), carried over into approximately the first half of the delay period (*p* = 0.006) and then increased again after verb onset (0.44 to 2.35 s, *p* = 0.006; unit-level LME cluster-based permutation tests, n = 77 units, all *p* values were Bonferroni corrected across the three periods; Fig. 3d). Because the images were absent from the screen throughout the delay and verb periods, this elevated FR during the verb period reflects internally generated reinstatement of stimulus-specific information rather than a sensory response. To ensure that verb-period reinstatement was not a mere continuation of persistent delay activity, we compared firing rates on preferred trials during the verb period with that during the late delay period (the 1 s preceding verb onset). This analysis yielded a *period* (delay, verb) × *image* (preferred, non-preferred) interaction (*p* = 0.036; Fig. 3d, inset), confirming enhanced FR of image-tuned units after verb onset.

Examining the role of these image-tuned units in successful vs. unsuccessful encoding of their preferred image, no subsequent-memory cluster survived correction across the three periods (*p*s > 0.108; Fig. 3e). Likewise, the corresponding period-averaged analysis showed no significant period × memory interaction (*p*s > 0.132; Fig. S3). In sum, image-specific reinstatement occurred during the verb period, largely irrespective of later memory outcome. Instead, the critical factor predicting subsequent memory was the extent of hippocampal engagement coinciding with this increase in FR of image-tuned units.

### Hippocampal HFA correlates with neural image reinstatement during successful binding

Increases in hippocampal HFA and neural reinstatement of the image therefore spanned the same time window during successful encoding. If hippocampal HFA draws upon stimulus representations to retroactively facilitate verb-image binding, the two signals should covary from trial to trial, and should do so selectively when the association is later remembered. For each image-tuned unit and hippocampal contact in the same participant, we paired the spike count over the verb period ([0, RT]) with the mean hippocampal HFA recorded over the same window on the same trial (Fig. 4a, b). Then we fit the HFA slope (HFA-FR correlation) on preferred/non-preferred × remembered/forgotten trials using a trial-level Poisson generalized linear mixed model (GLMM) that accounted for the trial-specific length of the verb period, so that the estimated slopes describe firing rate. To minimise potential contributions from local neuronal activity to the HFA estimate within the hippocampus, the primary analysis used only non-hippocampal image-tuned units (n = 54).

**Figure 4.**
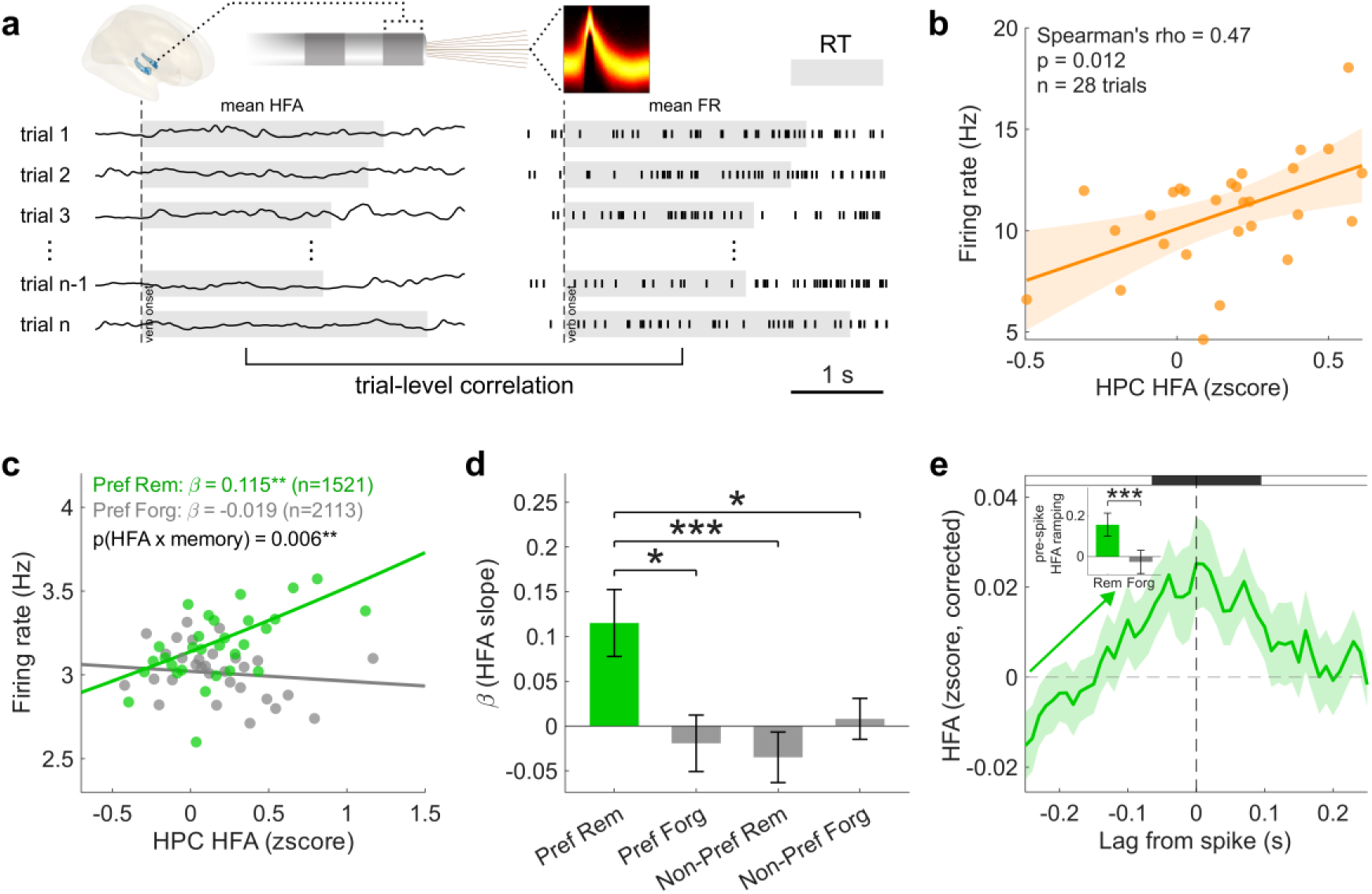
Trial-by-trial coupling between hippocampal HFA and FR of image-tuned neurons during the verb period. **a)** Analysis schematic. For each image-tuned unit paired with a hippocampal contact from the same participant, mean hippocampal HFA and unit firing rate were quantified over the verb period ([0, RT]) on preferred-image trials, and their trial-by-trial relationship was assessed. **b)** Example unit–contact pair. Trial-level mean firing rate ([0, RT], Hz) is plotted against trial-level mean hippocampal HFA ([0, RT], z-score; n = 28 preferred trials). Each dot represents one trial; the line and shading indicate the linear fit and its 95% confidence interval. **c)** Trial-level relationship between hippocampal HFA and firing rate in non-hippocampal image-tuned units (n = 54 from 10 participants). Lines show fitted slopes for preferred remembered and preferred forgotten trials from a trial-level Poisson generalized linear mixed model (GLMM, Poisson with log link: spike count∼ HFA × image × memory + (1|subject) + (1|subject:trial) + (1|subject:region:unit:hpc_contact); slopes are plotted on the firing-rate scale). Dots represent trials binned by HFA for visualization only. \*\**p* < 0.01. **d)** Estimated HFA effects (β ± s.e.) for preferred R, preferred F, non-preferred R and non-preferred F trials from the same Poisson GLMM as in (c). Brackets show post hoc contrasts of preferred R against each of the other three trial types, Bonferroni-corrected across the three comparisons (preferred forgotten *p* = 0.017; non-preferred remembered *p* < 0.001; non-preferred forgotten *p* = 0.041). \**p* < 0.05, \*\*\**p* < 0.001. **e)** Spike-locked hippocampal HFA during the verb period in non-hippocampal image-tuned units (n = 54, preferred R trials). Band-filtered HFA (50:10:150Hz) was corrected by subtracting the mean of trial-shuffled surrogate data (200 draws). Line and shading indicate mean ± s.e.m. across units. Horizontal black bar indicates significant time points where HFA exceeds the surrogates (one-sided unit-level LME cluster-based permutation, HFA ∼ condition + (1|subject) + (1|subject:region:unit)). Inset, pre-spike HFA ramp quantified as the fitted HFA slope over [−0.25, −0.10] s in preferred R and F trials (LME: HFA ∼ time × memory + (1|subject) + (time|subject:region:unit)). The upper bound of −0.10 s was chosen to avoid contamination from spike-triggered HFA leakage. \*\*\**p* < 0.001.

The GLMM yielded a significant three-way interaction: the modulation of the HFA-FR association by image preference differed as a function of subsequent memory (HFA × image × memory: beta = 0.177, s.e. = 0.048, *p* < 0.001). Specifically, within R trials, HFA tracked FR more strongly for preferred than non-preferred images (HFA × image: beta = 0.150, s.e. = 0.035, *p* < 0.001; Fig. 4c), whereas within F trials it did not (beta = −0.027, s.e. = 0.033, *p* = 0.406). Further unpacking the four cells (preferred/non-preferred × R/F), the HFA-FR slope was positive and significant only on *preferred R* trials (beta = 0.115, s.e. = 0.037, *p* = 0.002), and the *preferred R* slope exceeded each of the other three (*preferred F*, *non-preferred R*, and *non-preferred F*; *p*s ≤ 0.041, Bonferroni-corrected; Fig. 4d). The same pattern was obtained when all 77 image-tuned (including hippocampal) units were included (HFA × memory interaction: beta = 0.098, s.e. = 0.036, *p* = 0.007; Fig. S4). Hippocampal HFA thus selectively correlated with stimulus-specific firing when the verb-image association was successfully encoded.

### Hippocampal HFA increase precedes image reinstatement

Does hippocampal HFA during successful binding precede or follow firing of stimulus-selective units? To address this question, we computed spike-locked hippocampal HFA on preferred R trials (using a band-filtered HFA estimate with higher temporal precision than the multitaper estimate used above) and corrected each unit’s peri-spike time series by subtracting trial-shuffled surrogates from the same unit (200 draws; n = 54 non-hippocampal image-tuned units). Hippocampal HFA around the spikes of image-tuned units exceeded the surrogate distribution (*p* = 0.008, LME cluster-permutation test; Fig. 4e), corroborating the trial-specific coupling between these two signals. Critically, hippocampal HFA already increased before the spikes of image-tuned units: fitting HFA against time over the pre-spike window ([−0.25, −0.10] s), chosen to avoid contamination by filter-induced leakage from the spike itself (Fig. S5a), revealed a significant positive slope (beta = 0.153, s.e. = 0.057, *p* = 0.009; Fig. 4e, inset). Importantly, this pre-spike increase in hippocampal HFA was specific to later-remembered associations. Applying the same analysis to *preferred forgotten* trials, with surrogates drawn from each unit’s own *preferred forgotten* trials, revealed no significant hippocampal HFA cluster (ps > 0.152; Fig. S5) and no pre-spike increase (beta = −0.026, s.e. = 0.057, *p* = 0.644). Moreover, the hippocampal HFA slope was significantly steeper on R than on F trials (time × memory interaction, beta = 0.179, s.e. = 0.044, *p* < 0.001; Fig. 4e, inset). Hippocampal HFA increases thus preceded firing of image-tuned units, and this was only the case when the verb-image association was later remembered. This temporal ordering is consistent with the hippocampus retroactively recruiting stimulus representations in service of associative binding.

## Discussion

How does the brain form a single memory from two events that are experienced separately in time? By resolving hippocampal activity and stimulus-specific neuronal firing within the same encoding episodes (Fig. 1), our findings suggest a retroactive solution to this problem. Hippocampal HFA predicted subsequent associative memory only when the second event arrived, rather than during initial processing of the first event or the intervening delay (Fig. 2). At this same moment, neurons selective for the preceding image reinstated stimulus-specific firing despite the image no longer being visually present (Fig. 3). Critically, successful encoding was characterised not primarily by stronger reinstatement itself, but by the selective coupling of this reinstatement to hippocampal HFA (Fig. 4c). Hippocampal activity furthermore increased before spikes of stimulus-selective units on later-remembered trials (Fig. 4e). Together, these findings suggest that binding across time does not require continuous hippocampal engagement across the temporal gap. Instead, when new information arrives, the hippocampus may transiently recruit a representation of the preceding event, bringing past and present information into a common window for associative encoding.

Previous human neuroimaging work has implicated the hippocampus and neighbouring medial temporal regions in forming associations between temporally discontinuous events (Hales & Brewer, 2011; Hales et al., 2009; Qin et al., 2007; Staresina & Davachi, 2009), but the sluggishness of the BOLD response precluded determining when within such an episode the critical hippocampal operation occurs. Here, the hippocampal subsequent-memory effect emerged only after presentation of the verb, when both event elements had been experienced, and was absent during the preceding image and delay periods. This does not imply that the hippocampus cannot maintain information across short delays. In fact, human intracranial studies have demonstrated persistent hippocampal activity during working-memory maintenance, including activity that predicts later long-term memory (Axmacher et al., 2010; Daume et al., 2024). Rather, in the present associative-learning setting, successful binding was not indexed by sustained hippocampal engagement across the gap but by engagement when the later information became available.

This temporal profile converges with recent causal evidence from trace conditioning in mice (Puhger et al., 2024). That study found little evidence for sustained CA1 activation during the stimulus-free trace interval, whereas CA1 activity increased strongly after the later aversive stimulus. Critically, silencing CA1 immediately after this event impaired subsequent learning.

Our results show that the same temporal principle applies to intentional associative/episodic memory rather than only fear conditioning. Moreover, our data suggest that hippocampal engagement at the second event does not just signal successful encoding in general, but coincides with the return of information in image-tuned units that must be linked to the present.

Regarding the dynamics of image-tuned units, we observed that stimulus-selective firing weakened substantially after the image disappeared and increased again ∼500 ms after verb onset (Fig. 3d). This argues against a simple maintenance account in which the first event remains continuously represented until the second arrives. At first sight, the return of stimulus information resembles proposals of ‘activity-silent’ working memory, in which latent representations can be recovered by perturbing the network with a simple, non-specific visual impulse (Stokes, 2015; Wolff et al., 2017). However, the present reinstatement seems more consistent with a mnemonic retrieval operation than with a generic working-memory ping (here, the verb onset). Unlike the fast (∼200 ms) pinging response, reinstatement emerged only after several hundred milliseconds, and closely resembles delayed concept-cell reactivation when a current stimulus or linguistic cue refers back to a preceding concept (Bausch et al., 2021; Dijksterhuis et al., 2024; Staresina et al., 2019).

Importantly, reinstatement alone was insufficient for successful memory encoding. Image-tuned firing returned during the verb period largely irrespective of whether the association was subsequently remembered (Fig. 3e). Instead, what differentiated successful encoding was the relationship between this content-specific firing and hippocampal activity. Hippocampal HFA tracked firing selectively when a unit’s preferred image was the relevant preceding event and the association was later remembered (Fig 4c-d). This interaction suggests that stimulus-selective units specify what information is currently being reinstated, whereas hippocampal engagement may coordinate its effective mnemonic binding with the word. This is consistent with the long-held notion that the hippocampus serves as an indexing or relational system that orchestrates distributed representations rather than storing all constituent content itself (Eichenbaum et al., 1992; McClelland et al., 1995; Teyler & DiScenna, 1986).

The spike-locked analysis (Fig 4e) further revealed that on successfully encoded trials, hippocampal HFA increased before spikes in non-hippocampal image-tuned units. This ordering is consistent with hippocampal engagement facilitating recruitment of the preceding representation rather than merely reflecting its arrival. Nevertheless, temporal precedence does not establish causal direction, and establishing a causal role will require perturbation, e.g., interference with hippocampal activity around the moment when the second event arrives.

Retroactive recruitment in service of memory formation may also extend beyond the deliberate associative learning examined here. In our task, participants knew that the incoming verb had to be related to the preceding image (which likely also explains the stimulus-specific reinstatement irrespective of later memory success, Fig. 3e). In everyday memory, however, the relevance of an earlier experience sometimes becomes apparent only later and without any prior intention to associate the events. For instance, a later salient experience can selectively strengthen memory for earlier, initially inconsequential but conceptually related information (Dunsmoor et al., 2015). Recent neuroimaging work suggests that such retroactive enhancement is accompanied by reactivation of preceding event representations in the hippocampus and stimulus-relevant neocortex, together with increased hippocampal– neocortical coupling during subsequent emotional learning (Zhu et al., 2022). These phenomena unfold over different timescales and probably involve additional mechanisms, including consolidation. Nonetheless, they suggest a potentially shared principle: when present information changes the relevance of the recent past, hippocampal engagement may promote reinstatement of that past so that a memory can be formed or updated retroactively.

There are several open questions for future work. The content space was deliberately constrained to four repeatedly encountered images, allowing stimulus-specific neurons to be identified independently. Whether the same mechanism operates for richer, one-shot representations of naturalistic experience remains to be established. Moreover, although intracranial recordings provide exceptional temporal precision, the data are necessarily obtained from patients with epilepsy and from clinically determined recording locations. As such, our neuronal recordings were restricted to the medial temporal lobe and it would be important to understand how the hippocampus interacts with fronto-parietal executive and attention systems upon encountering the second event.

Taken together, our results identify a highly specific profile of hippocampal involvement in binding events across time. Rather than carrying the first event continuously across the intervening gap, successful encoding was associated with hippocampal engagement when the second event arrived, accompanied by reinstatement of the earlier event’s neuronal representation and by selective coupling between the two signals. The hippocampus may therefore bridge temporal discontinuities by bringing the past back into the present, creating a transient window in which information separated in physical time can nevertheless be bound into a common memory.

## Methods

### Participants

We analysed intracranial EEG (iEEG) recordings from 15 patients with drug-resistant epilepsy undergoing invasive presurgical seizure monitoring at the Epilepsy Center Freiburg, Freiburg im Breisgau, Germany (40 ± 11 years, mean ± s.d.; 6 female, 9 male; table S1). All patients were evaluated for potential neurosurgical treatment and were implanted with depth electrodes for clinical purposes. 13 patients additionally had microwires allowing single-unit recordings. All participants gave written informed consent, and the study was approved by the ethics committee of the University Medical Center Freiburg, Germany.

### Experimental design

Each participant performed an image-verb associative memory task consisting of a localiser, an encoding and a retrieval period. In the localiser (160 trials), one of four images (river, church, piano, elephant) was presented for 1 s, with an inter-trial interval of 1 s plus a jitter of 0, 0.1, 0.2 or 0.3 s; participants performed a one-back task, pressing a button whenever an image was identical to the preceding one. Immediate stimulus repeats and trials containing a button press were excluded from all subsequent analyses, so that the localiser provided image-evoked responses uncontaminated by motor or repetition effects.

During encoding, each trial began with a fixation cross (2 s plus a jitter of 0, 0.1, 0.2 or 0.3 s), followed by one of the four images (1.5 s plus a jitter of 0, 0.1, 0.2 or 0.3 s), a blank delay (1.75 s plus a jitter of 0, 0.25, 0.5 or 0.75 s) and lastly a verb. Participants judged whether the image– verb pairing was plausible (left key, plausible; right key, implausible), responding at their own pace within a window of 2–5 s after verb onset (minimum 2 s on screen irrespective of the response); the mean plausibility reaction time (RT) was 2.5 ± 0.5 s. Encoding and retrieval alternated in blocks, so that each encoding block was followed by retrieval of the pairs studied in that block. Block length was adapted to each participant’s performance to keep the task at a comparable level of difficulty: a block comprised 16 image–verb pairs by default, and was lengthened to 32 pairs (2 participants) or shortened to 8 pairs (1 participant) for participants with high or low cued-recall accuracy, respectively. This adjustment was made after the initial practice block and was then held constant for the remainder of the session, yielding 4–12 blocks per participant (114 ± 13 trials per participant). Encoding and retrieval blocks were separated by a distractor task lasting at least 30 s, in which participants covertly counted colour changes of a central fixation cross. During retrieval, a fixation cross (2 s plus a jitter of 0, 0.1, 0.2 or 0.3 s) was followed by the verb cue (2–10 s; minimum 2 s on screen irrespective of the response), and participants indicated which of the four images had been paired with that verb by pressing one of four keys, each labelled with a letter identifying one image; of note, participants were not forced to guess, but could press a dedicated key to indicate they had forgotten the target image. A trial was scored as remembered if the image selected at retrieval matched the image encoded with that verb, and as forgotten otherwise.

### iEEG macro recording and preprocessing

iEEG data were acquired using a Compumedics EEG system (Compumedics, Abbotsford, Victoria, Australia) at a sampling rate of 2,000 Hz, with depth electrodes provided by Ad-Tech (Ad-Tech Medical Instrument Corp., Racine, WI, USA), and were referenced to Cz.

Preprocessing was performed in FieldTrip (Oostenveld et al., 2011). Continuous data were high-pass filtered at 0.1 Hz and band-stop filtered around line noise and its harmonics (49–51, 99–101, 149–151, 199–201 and 249–251 Hz), then resampled to 1,000 Hz. iEEG traces were visually inspected, and contacts with physical noise or located outside the brain were removed. Contacts were labelled as medial temporal lobe (MTL) contacts if they fell within the amygdala, hippocampus, parahippocampal cortex or rhinal (entorhinal/perirhinal) cortex, as determined visually in native space using LeGUI (Davis et al., 2021). Each MTL contact was then re-referenced against its nearest white-matter contact on the same depth electrode. Across the 15 participants this yielded 127 MTL contacts (amygdala, n = 29; hippocampus, n = 38; parahippocampal cortex, n = 19; rhinal cortex, n = 41). For group-level visualisation, contact coordinates were normalised to MNI space and displayed on a template brain in FieldTrip.

### iEEG high frequency activity (HFA)

Two complementary estimates of high-frequency activity (HFA) were computed, both epoched relative to image onset ([−0.5, 2] s), delay onset ([−0.5, 2] s) and verb onset ([−0.5, 3] s). The two differ in their trade-off between spectral resolution and temporal precision: the multitaper estimate is frequency-resolved but temporally smoothed by its 400-ms analysis window, whereas the band-pass/Hilbert estimate is spectrally coarser but temporally precise enough to resolve HFA around individual spikes.

For the time–frequency analyses and the main HFA time courses (Figs. 2 to 4 and fig. S1), spectral power was estimated with a multitaper approach (FieldTrip), using a Hanning taper and 5-cycle windows for low frequencies (2–29 Hz, 1-Hz steps), DPSS tapers ±10 Hz spectral smoothing and 400-ms windows for high frequencies (30–200 Hz, 5-Hz steps), and progressing in 10 ms steps. For each contact and frequency, power was epoched relative to image, delay and verb onset. The three epochs were then concatenated across trials to define the encoding period, and power was z-scored against the mean and standard deviation of that period, after removing across-trial outliers (MATLAB function *isoutlier*); the z-scored values were then averaged across the 50–150 Hz band to capture high-frequency activity (HFA). Period-wise HFA means were computed over [0, 1.5] s (image), [0, 1.75] s (delay) and [0, RT] (verb, using each trial’s plausibility RT).

For the spike-locked analysis, HFA was instead computed by band-pass filtering each contact into ten 10-Hz sub-bands spanning 50–150 Hz (fourth-order Butterworth), taking the squared magnitude of the Hilbert transform and downsampling the resulting signal to 100 Hz. Each sub-band was z-scored per contact after outlier removal as described above, and the z-scored sub-bands were averaged.

### Microwire recordings and spike sorting

Neuronal signals were recorded with Behnke–Fried depth electrodes (Ad-Tech Medical Instrument Corp., Racine, WI, USA), each carrying a bundle of nine platinum–iridium microwires (40 µm in diameter) protruding from the tip of the depth electrode. Eight microwires served as recording channels, from which local field potentials were later extracted, and the ninth served as a reference. Microwire signals were sampled at 30 kHz using the NeuroPort system (Blackrock Microsystems, Salt Lake City, UT, USA) and band-pass filtered at 0.3–7,500 Hz. Microwires were implanted in several MTL regions, including the amygdala, hippocampus, parahippocampal cortex, entorhinal cortex, and perirhinal cortex.

Spikes (action potentials) were detected and sorted across the entire recording using Combinato (Niediek et al., 2016). Sorting results were further curated by two experienced annotators, who removed artifacts and merged clusters on the basis of spike waveform shape, inter-spike-interval distributions, and cross-correlograms. Across all sessions, we identified 440 MTL units (276 single units and 164 multi-units). For each unit, per-trial firing rates were computed in the same task epochs as the macro data, using a bin width of 1 ms (1,000 Hz).

### Image-tuned unit detection

As mentioned above, 13 participants had microwire recordings. One was excluded because only three units could be isolated, leaving 12 participants with usable spike data; in one of these the localiser was unavailable, and encoding-trial responses to the images were used instead (see below). Only non-repeating, non-response localiser trials were used, and only units with a mean firing rate above 0.1 Hz across events were analyzed. Across the 12 participants, 440 units passed the 0.1 Hz firing-rate criterion and were assessed for image tuning. For each unit and trial, we computed a baseline firing rate ([−1, −0.05] s) and an image-induced firing rate ([0, 1] s). A unit’s preferred image was the image eliciting the highest mean induced firing rate. Units had to satisfy two criteria to be classified as *image-tuned* (Kunz et al., 2024). Criterion 1 tested whether induced firing was higher for the preferred than the non-preferred images using a one-sided two-sample t-test; the empirical t statistic had to exceed the 95th percentile of a null distribution from 1,000 permutations in which image labels were shuffled across trials (the preferred image being re-selected on each surrogate). Criterion 2 tested whether the baseline-corrected, time-resolved firing rate showed a sustained preferred > non-preferred response: firing rates were Gaussian-smoothed (0.1 s kernel), baseline-corrected, and compared over [0, 1] s with a one-sided cluster-based permutation test for independent samples (FieldTrip, indepsamplesT, maxsum cluster statistic, 1,000 randomisations, alpha = 0.05). A unit was classified as image-tuned only if both criteria were met. This yielded 77 of 440 units (17.5%), distributed across MTL regions and approximately balanced across the four preferred images.

Next, we epoched the firing rates of image-tuned units relative to image, delay and verb onset. The data were then smoothed with a 0.4 s Gaussian kernel (MATLAB function *smoothdata*), matching the temporal smoothing of the HFA estimates, and normalised to the pre-image baseline ([−1, −0.2] s). Note that firing rates were normalised differently from HFA, which was z-scored against the whole encoding period, but in both cases a single reference per contact or unit was applied to all three trial phases, so comparisons across phases and conditions are unaffected.

For the aforementioned participant for whom encoding trials were used to detect image-tuned units, the data were taken from the image epoch of the encoding task, using the same induced ([0, 1] s) and baseline ([−1, −0.05] s) windows and the identical two-criterion pipeline applied to the localiser in all other participants. Because all reinstatement and coupling analyses concern the verb period, which begins at least 3.25 s after image onset, the window used to define a unit’s preferred image and the window used to test it do not overlap. Excluding the participant altogether left the trial-level HFA–firing association unchanged (HFA × image × memory: beta = 0.189, s.e. = 0.048, *p* < 0.001).

### Linear mixed-effects (LME) tests

Group-level HFA and firing-rate time courses were tested with cluster-based permutation tests combined with linear mixed-effects statistics to account for the hierarchical structure of our data (FieldTrip with a custom LME statistic function; Aarts et al., 2014). All tests were two-tailed unless otherwise specified and reported cluster *p* values are additionally Bonferroni-corrected across the three encoding trial periods (image, delay and verb), that is, multiplied by three. For HFA analyses (Fig. 2 and figs. S1 and S2), trials were first averaged within condition (R, F) separately for each contact, so that one observation is one time point in one condition at one contact. The later-remembered versus later-forgotten contrast was then modelled as:

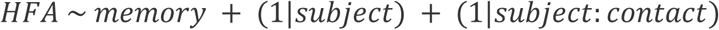

where HFA is the z-scored 50–150 Hz power of one contact at a given time point, memory is a categorical fixed effect coding subsequently remembered versus forgotten trials, subject is the participant identifier and contact is the contact identifier nested within participant. The time-resolved LME was run with 1,000 permutations and a cluster alpha of 0.05; the same framework was applied in two dimensions (frequency × time) for the time–frequency contrast (fig. S1).

For single-/multi-unit time courses (Fig. 3), trials were likewise averaged within condition separately for each unit, so that one observation is one time point in one condition at one unit. Firing rates were smoothed with a 400-ms Gaussian kernel to match the temporal smoothing of the HFA estimates, and the contrast of interest (preferred versus non-preferred, or remembered versus forgotten) was modelled as

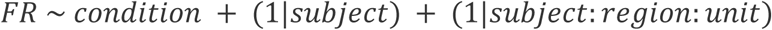

where FR is the smoothed firing rate of one unit at a given time point, condition is a categorical fixed effect coding the contrast of interest, subject is the participant identifier, region is the MTL region in which the unit was recorded and unit is the unit identifier nested within participant and region.

The effects of period (image, delay, verb) × memory (R, F) interaction were tested with 2 × 2 LMEs of the form *HFA* ∼ *period* ∗ *condition* + (1|*subject*) + (1|*subject*: *c*ℎ*annel*) for macro data and *FR* ∼ *period* ∗ *condition* + (1|*subject*) + (1|*subject*: *region*: *unit*) for units; the interaction term is reported. The effect of region for R-F difference was tested using an LME model (*HFA* ∼ *region* + (1|*subject*)) for HFA data. Post-hoc contrasts from the same model tested (i) the R − F difference against zero in each region (Bonferroni corrected), and (ii) the hippocampus against each of these regions (Bonferroni corrected).

Unless stated otherwise, error bars and shaded bands denote the s.e.m. across the relevant observation unit (contacts or units), and significance is reported as \**p* < 0.05, \*\**p* < 0.01, \*\*\**p* < 0.001; n.s., *p* > 0.05.

### Trial-by-trial coupling between HFA and image-tuned firing

To test whether hippocampal HFA correlates with image-tuned firing across trials, each image-tuned unit was paired with every hippocampal contact from the same participant. For each trial we computed, over the verb window [0, RT], the unit’s spike count and the contact’s mean z-scored hippocampal HFA (50–150 Hz).

Pooling across pairs, we fitted a trial-level Poisson generalised linear mixed model (GLMM; log link, Laplace approximation) with the spike count as the response and log(RT) as an offset, to account for the variable window duration across trials:

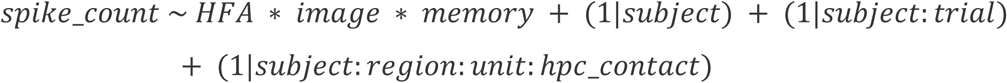

where image coded preferred versus non-preferred trials and memory coded remembered versus forgotten trials. The (1|subject) random effect accounts for subject-level variance, (1|subject:trial) accounts for trial-level variance and (1|subject:region:unit:hpc_contact) accounts for pair-level variance. The HFA slope in each of the four image × memory cells, and the differences between the preferred-remembered slope and each of the other three cells, were obtained as post hoc contrasts of the three-way GLMM. We focus on the non-hippocampal (non-HPC) units only to rule out spurious within-hippocampus firing-to-HFA associations.

### Spike-locked hippocampal HFA

To assess the temporal relationship between hippocampal HFA and image-tuned spikes, we computed spike-locked HFA separately on preferred-R and preferred-F trials. Per participant, we used the band-filtered, z-scored hippocampal HFA (50–150 Hz sub-bands, described above), because this estimate is temporally more precise than the multitaper estimate (400-ms smoothing). For every spike emitted in the verb window [0, RT] of a trial of the given condition, the HFA time course was sampled at lags from −0.25 to 0.25 s (100 Hz, i.e. 10-ms steps). A trial-shuffle surrogate was constructed by sampling the same spike time and lags from a randomly chosen trial of the same condition and the same unit (200 draws), and the corrected peri-spike HFA was the raw curve minus the mean surrogate. Analyses were run on non-HPC units (n = 54 in both conditions).

For each condition, over the lag window [−0.25, 0.25] s, the real and surrogate curves were compared with a unit-level LME cluster-based permutation test ( *HFA* ∼ *condition* + (1|*subject*) + (1|*subject*: *region*: *unit*), with condition labels (real versus shuffled) permuted within unit, 1,000 permutations).

As a next step, we tested the pre-spike HFA slope. The HFA filter was applied in two passes (zero-phase), which is acausal, so power following a spike can in principle leak backwards in time. We measured that leakage rather than assuming it to be negligible, by passing a unit impulse through the identical filter path. 97.7% of the filter’s total response fell within +/-100 ms of the impulse, and 1.1% fell inside the window [−0.25, −0.10] s (Fig. S5a). We then chose the window to limit contamination from post-spike HFA leakage. Pre-spike slope was quantified on the corrected curves over [−0.25, −0.10] s by fitting an interaction model, *HFA* ∼ *time* ∗ *memory* + (1|*subject*) + (*time*|*subject*: *region*: *unit*), with R trials as the reference level. The time × memory interaction tested whether the slope differed between R and F trials, the (time|subject:region:unit) random slope allowed the pre-spike HFA slope to vary across units.

## Data and code availability

The data and code supporting the findings of this study will be made publicly available through the German Neuroinformatics Node (G-Node; https://gin.g-node.org/) upon publication.

## Acknowledgements

We thank Tim Guth for helpful discussions on data analyses.

## Author contributions

MP and BPS designed the study. MFK, AB, and LK conducted the experiments. XH pre-processed the data with input from MSK. XH and BPS planned the analyses. XH analysed the data with input from MSK and BPS. XH and BPS designed and prepared the figures. LK, ASB and BPS supervised and administered the project and secured funding. XH and BPS wrote the manuscript. All authors discussed the results and commented on the manuscript.

## Supplementary

**Table S1.** Number of intracranial contacts and units across participants.

| patient | macro_amy | macro_hpc | macro_phc | macro_rhinal | units | img_tuned_units | img_tuned_nonHPC_units |
| --- | --- | --- | --- | --- | --- | --- | --- |
| p01 | 2 | 2 | 1 | 2 | 54 | 3 | 3 |
| p02 | 2 | 2 | 0 | 2 | 27 | 2 | 1 |
| p03 | 4 | 4 | 1 | 7 | 16 | 0 | 0 |
| p04 | 0 | 2 | 1 | 2 | 13 | 1 | 1 |
| p05 | 1 | 4 | 2 | 2 | 65 | 8 | 3 |
| p06 | 2 | 2 | 1 | 1 | / | / | / |
| p07 | 2 | 2 | 2 | 2 | 46 | 19 | 14 |
| p08 | 2 | 2 | 3 | 2 | 33 | 4 | 3 |
| p09 | 0 | 4 | 2 | 1 | 67 | 20 | 15 |
| p10 | 3 | 5 | 0 | 9 | 42 | 7 | 4 |
| p11 | 2 | 2 | 1 | 1 | / | / | / |
| p12 | 2 | 2 | 2 | 2 | 19 | 2 | 1 |
| p13 | 2 | 1 | 1 | 3 | 33 | 9 | 9 |
| p14 | 1 | 2 | 0 | 3 | / | / | / |
| p15 | 4 | 2 | 2 | 2 | 25 | 2 | 0 |
| Total | 29 | 38 | 19 | 41 | 440 | 77 | 54 |
| Mean±SD | 1.9±1.2 | 2.5±1.1 | 1.3±0.9 | 2.7±2.3 | 36.7±18.3 | 6.4±6.7 | 4.5±5.3 |

**Figure S1.**
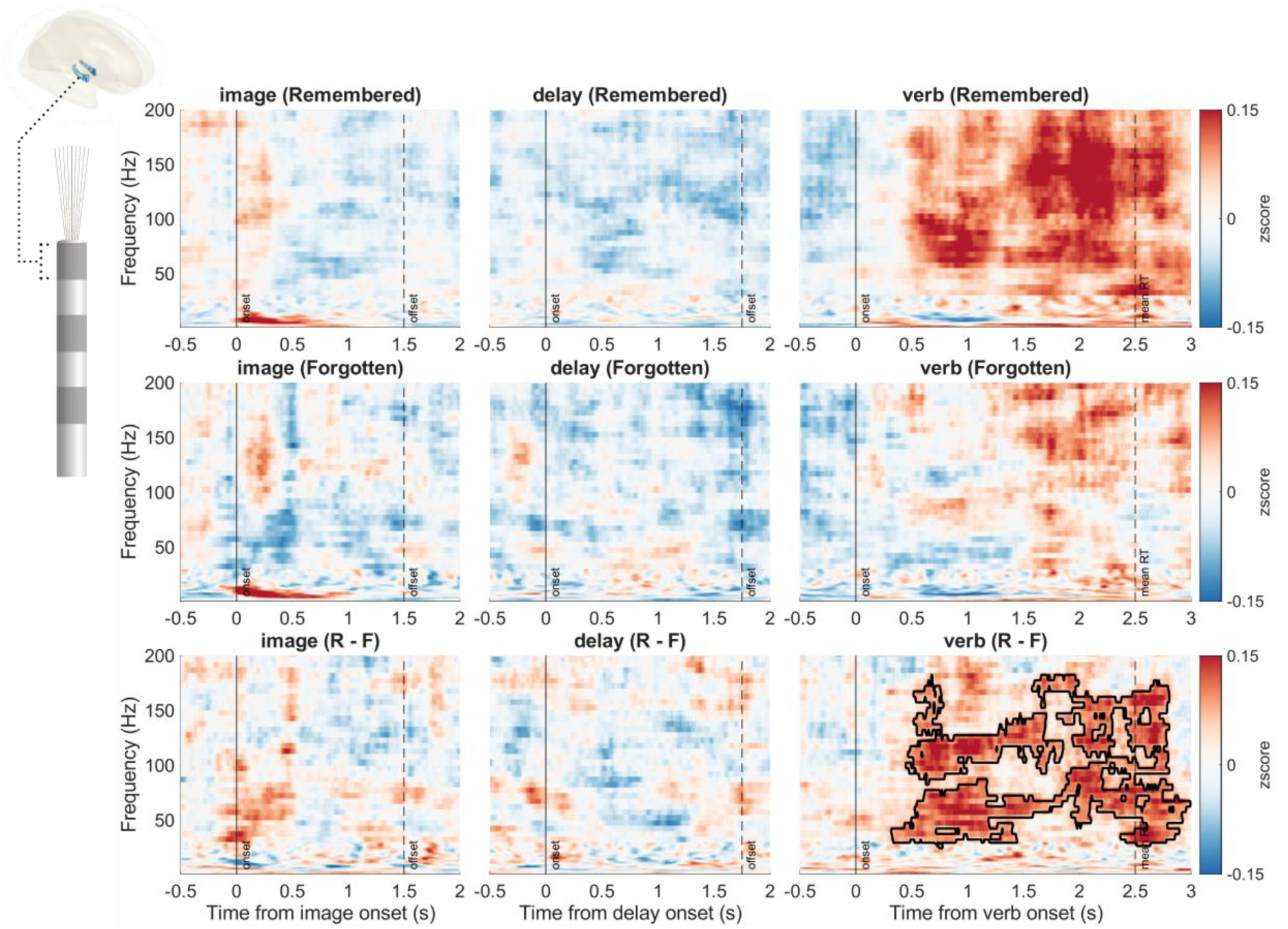
Hippocampal time–frequency subsequent-memory effect. Time–frequency representations (2–200 Hz; z-scored relative to the entire encoding period) aligned to image, delay and verb onset across hippocampal contacts (n = 38). Rows show later remembered trials (R, top), later forgotten trials (F, middle) and the R-minus-F difference (bottom). The black contour marks a significant R > F cluster (LME cluster-based permutation, power z-score ∼ memory + (1|subject) + (1|subject:contact), 1,000 permutations, *p* < 0.05).

**Figure S2.**
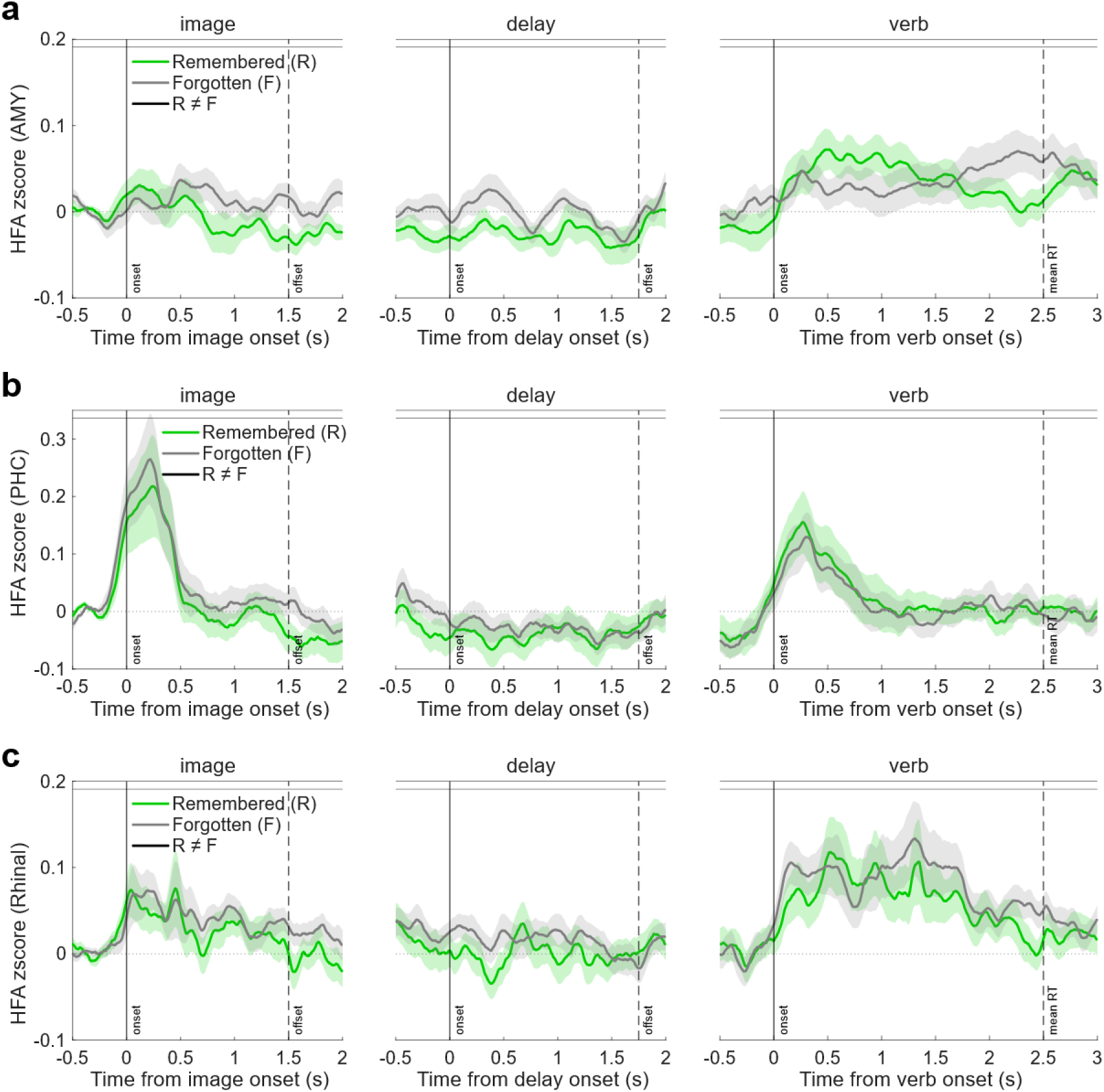
No subsequent memory effect in other MTL regions. HFA time courses (50–150 Hz; z-scored relative to the entire encoding period) for later remembered (R) and forgotten (F) trials, aligned to image, delay and verb onset, in (a) amygdala (n=29); (b) parahippocampal cortex (n=19); and (c) rhinal cortex (n=41). Lines and shading indicate mean ± s.e.m. across contacts. No cluster survived correction in any region or period (HFA ∼ memory + (1|subject) + (1|subject:contact), 1,000 permutations, Bonferroni-corrected across the image, delay and verb periods, *p* > 0.05).

**Figure S3.**
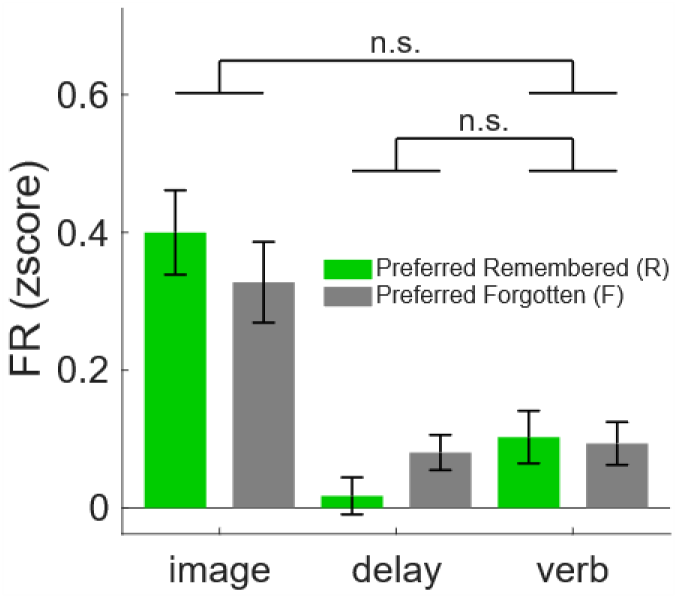
No significant period x memory interaction in image-tuned firing. Mean firing rate of image-tuned units for preferred later remembered and forgotten trials during the image ([0, 1.5] s), delay ([0, 1.75] s) and verb ([0, RT]) periods. Bars indicate mean ± s.e.m. across image-tuned units (n = 77). Period × memory interactions were assessed using LME models (FR ∼ period × memory + (1|subject) + (1|subject:region:unit)). n.s. *p* > 0.05.

**Figure S4.**
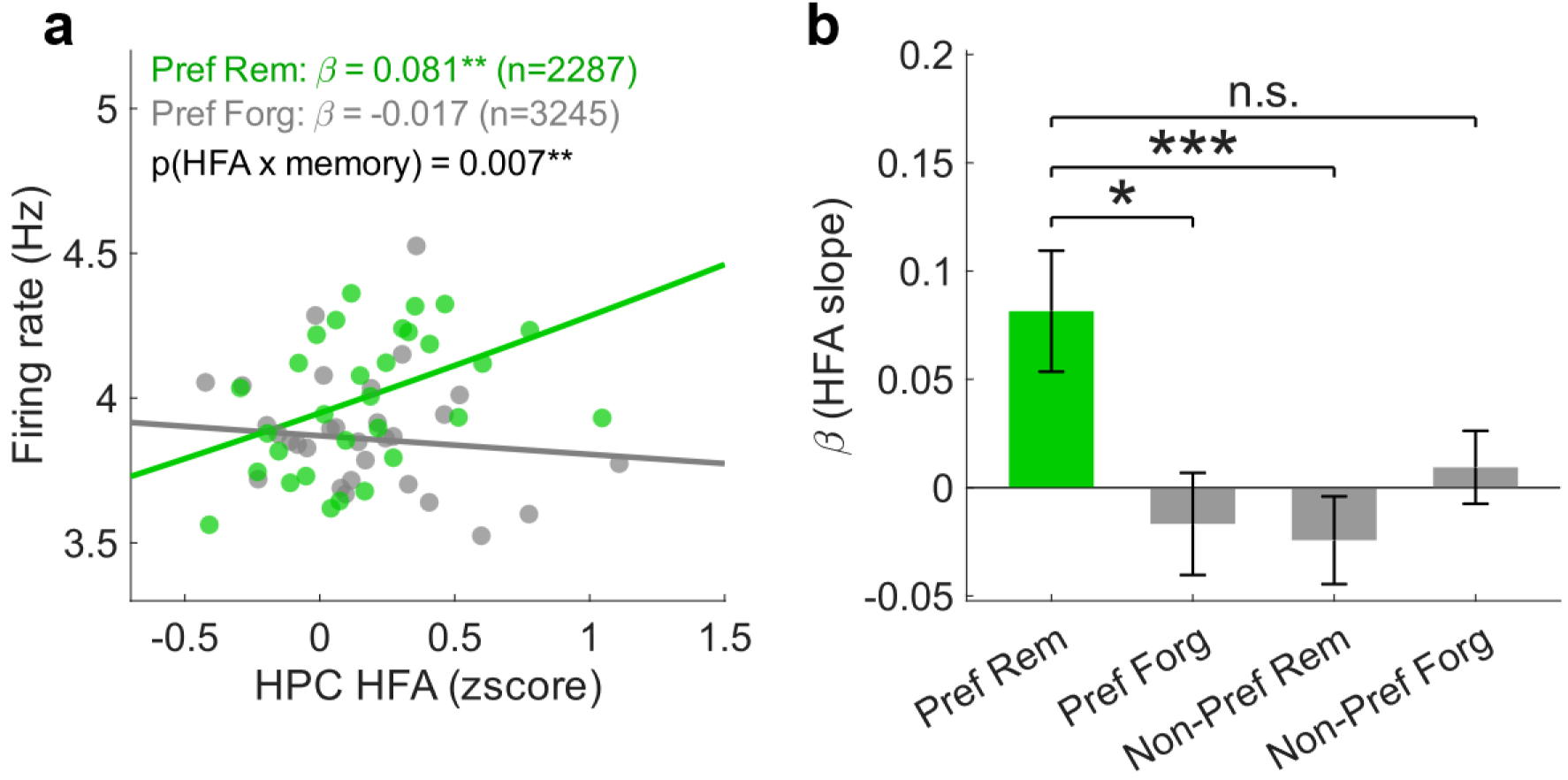
Trial-by-trial coupling between hippocampal HFA and image-tuned firing across all image-tuned units. **a)** Trial-level relationship between mean firing rate and hippocampal HFA during the verb period ([0, RT]) across all image-tuned units (including hippocampal units). Lines show fitted slopes for preferred remembered and preferred forgotten trials from a trial-level Poisson GLMM (GLMM, Poisson with log link: spike count∼ HFA × image × memory + (1|subject) + (1|subject:trial) + (1|subject:region:unit:hpc_contact), with log(RT) as an offset; slopes are plotted on the firing-rate scale). Dots represent trials binned by HFA for visualization only. \*\**p* < 0.01. **b)** Estimated HFA effects (β ± s.e.) for preferred later remembered, preferred later forgotten, non-preferred later remembered and non-preferred later forgotten trials from the same Poisson GLMM as in (a). \*\*\**p* < 0.001.

**Figure S5.**
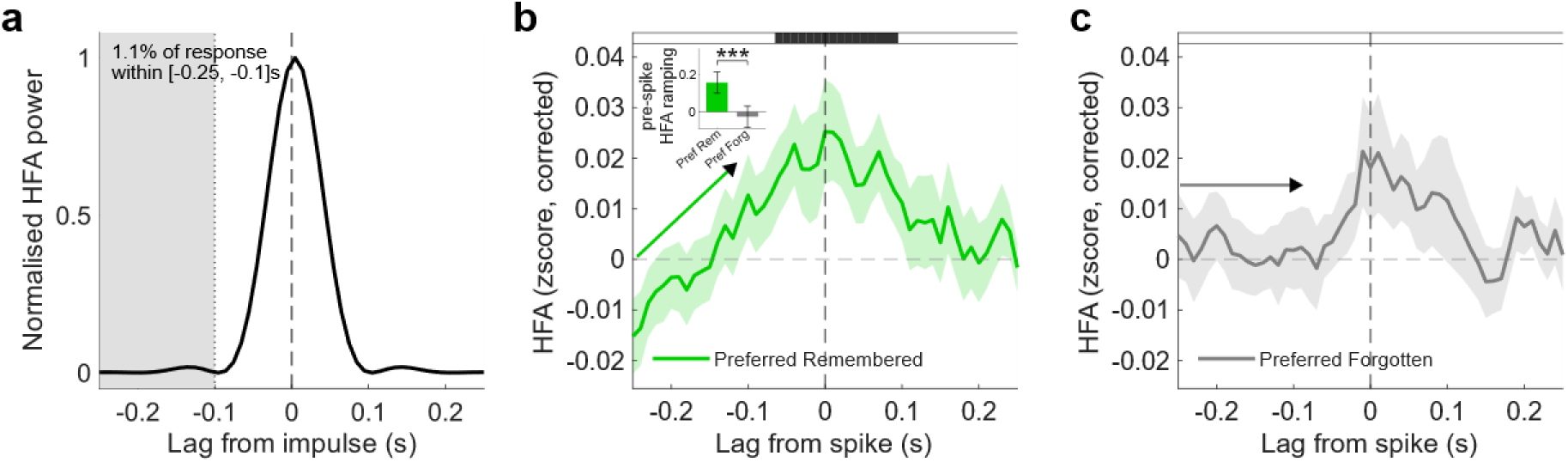
Spike-locked hippocampal HFA. **a)** Acausal spread of the HFA band-pass filters. Normalised impulse response of the two-pass (zero-phase) fourth-order Butterworth sub-band filters used for the spike-locked HFA estimate, averaged over the ten 10-Hz sub-bands and plotted on the spike-locked lag axis. 97.7% of the response falls within +/-100 ms of the impulse (dotted line indicates −0.1s), while the shaded pre-spike ramp window [−0.25, −0.10] s contains 1.1% of it. **b)** Spike-locked hippocampal HFA during the verb period in non-hippocampal image-tuned units (n = 54, preferred later remembered trials), same as Fig. 4e. Band-filtered HFA (50:10:150Hz) was corrected by subtracting the mean of trial-shuffled surrogate data (200 draws). Line and shading indicate mean ± s.e.m. across units. Horizontal black bar indicates significant lags where HFA exceeds the surrogate (one-sided unit-level LME cluster-based permutation, HFA ∼ condition + (1|subject) + (1|subject:region:unit)). Inset, pre-spike HFA ramp quantified as the fitted HFA slope over [−0.25, −0.10]s in preferred later remembered and forgotten trials (LME: HFA ∼ lag × memory + (1|subject) + (lag|subject:region:unit)). The upper bound of −0.10 s was chosen to avoid contamination from spike-triggered HFA leakage. \*\*\**p* < 0.001. **c)** Same as in (b) but for preferred later forgotten trials.

## References

Aarts, E., Verhage, M., Veenvliet, J. V., Dolan, C. V., & van der Sluis, S. (2014). A solution to dependency: Using multilevel analysis to accommodate nested data. Nature Neuroscience, 17(4), 491–496. 10.1038/nn.3648

Axmacher, N., Henseler, M. M., Jensen, O., Weinreich, I., Elger, C. E., & Fell, J. (2010). Cross-frequency coupling supports multi-item working memory in the human hippocampus. Proceedings of the National Academy of Sciences, 107(7), 3228–3233. 10.1073/pnas.0911531107

Bausch, M., Niediek, J., Reber, T. P., Mackay, S., Boström, J., Elger, C. E., & Mormann, F. (2021). Concept neurons in the human medial temporal lobe flexibly represent abstract relations between concepts. Nature Communications, 12, 6164. 10.1038/s41467-021-26327-3

Burke, J. F., Long, N. M., Zaghloul, K. A., Sharan, A. D., Sperling, M. R., & Kahana, M. J. (2014). Human intracranial high-frequency activity maps episodic memory formation in space and time. NeuroImage, 85 Pt 2(0 2), 834–843. 10.1016/j.neuroimage.2013.06.067

Clark, R. E., & Squire, L. R. (1998). Classical conditioning and brain systems: The role of awareness. Science, 280(5360), 77–81. 10.1126/science.280.5360.77

Clewett, D., DuBrow, S., & Davachi, L. (2019). Transcending time in the brain: How event memories are constructed from experience. Hippocampus, 29(3), 162–183. 10.1002/hipo.23074

Dan, Y., & Poo, M. (2004). Spike timing-dependent plasticity of neural circuits. Neuron, 44(1), 23–30. 10.1016/j.neuron.2004.09.007

Daume, J., Kamiński, J., Salimpour, Y., Gómez Palacio Schjetnan, A., Anderson, W. S., Valiante, T. A., Mamelak, A. N., & Rutishauser, U. (2024). Persistent activity during working memory maintenance predicts long-term memory formation in the human hippocampus. Neuron, 112(23), 3957–3968.e3. 10.1016/j.neuron.2024.09.013

Davachi, L. (2006). Item, context and relational episodic encoding in humans. Current Opinion in Neurobiology, 16(6), 693–700. 10.1016/j.conb.2006.10.012

Davis, T. S., Caston, R. M., Philip, B., Charlebois, C. M., Anderson, D. N., Weaver, K. E., Smith, E. H., & Rolston, J. D. (2021). LeGUI: A fast and accurate graphical user interface for automated detection and anatomical localization of intracranial electrodes. Frontiers in Neuroscience, 15, 769872. 10.3389/fnins.2021.769872

Dijksterhuis, D. E., Self, M. W., Possel, J. K., Peters, J. C., van Straaten, E. C. W., Idema, S., Baaijen, J. C., van der Salm, S. M. A., Aarnoutse, E. J., van Klink, N. C. E., van Eijsden, P., Hanslmayr, S., Chelvarajah, R., Roux, F., Kolibius, L. D., Sawlani, V., Rollings, D. T., Dehaene, S., & Roelfsema, P. R. (2024). Pronouns reactivate conceptual representations in human hippocampal neurons. Science, 385(6716), 1478–1484. 10.1126/science.adr2813

Dunsmoor, J. E., Murty, V. P., Davachi, L., & Phelps, E. A. (2015). Emotional learning selectively and retroactively strengthens memories for related events. Nature, 520(7547), 345–348.

Eichenbaum, H. (2004). Hippocampus: cognitive processes and neural representations that underlie declarative memory. Neuron, 44(1), 109–120. 10.1016/j.neuron.2004.08.028

Eichenbaum, H., Otto, T., & Cohen, N. J. (1992). The hippocampus—What does it do? Behavioral and Neural Biology, 57(1), 2–36. 10.1016/0163-1047(92)90724-I

Feldman, D. E. (2012). The spike-timing dependence of plasticity. Neuron, 75(4), 556–571. 10.1016/j.neuron.2012.08.001

Gelbard-Sagiv, H., Mukamel, R., Harel, M., Malach, R., & Fried, I. (2008). Internally generated reactivation of single neurons in human hippocampus during free recall. Science, 322(5898), 96–101.

Hales, J. B., & Brewer, J. B. (2011). The timing of associative memory formation: Frontal lobe and anterior medial temporal lobe activity at associative binding predicts memory. Journal of Neurophysiology, 105(4), 1454–1463. 10.1152/jn.00902.2010

Hales, J. B., Israel, S. L., Swann, N. C., & Brewer, J. B. (2009). Dissociation of frontal and medial temporal lobe activity in maintenance and binding of sequentially presented paired associates. Journal of Cognitive Neuroscience, 21(7), 1244–1254.

Kehl, M. S., Reber, T. P., Borger, V., Surges, R., Mormann, F., & Staresina, B. P. (2026). Sleep ripples drive single-neuron reactivation for human memory consolidation (p. 2026.03.27.714528). bioRxiv. 10.64898/2026.03.27.714528

Kunz, L., Staresina, B. P., Reinacher, P. C., Brandt, A., Guth, T. A., Schulze-Bonhage, A., & Jacobs, J. (2024). Ripple-locked coactivity of stimulus-specific neurons and human associative memory. Nature Neuroscience, 27(3), 587–599. 10.1038/s41593-023-01550-x

Lisman, J. E. (1999). Relating hippocampal circuitry to function: Recall of memory sequences by reciprocal dentate-CA3 interactions. Neuron, 22(2), 233–242. 10.1016/s0896-6273(00)81085-5

Mackay, S., Reber, T. P., Bausch, M., Boström, J., Elger, C. E., & Mormann, F. (2024). Concept and location neurons in the human brain provide the ‘what’ and ‘where’ in memory formation. Nature Communications, 15(1), 7926. 10.1038/s41467-024-52295-5

McClelland, J. L., McNaughton, B. L., & O’Reilly, R. C. (1995). Why there are complementary learning systems in the hippocampus and neocortex: Insights from the successes and failures of connectionist models of learning and memory. Psychological Review, 102(3), 419–457. 10.1037/0033-295X.102.3.419

Moyer, J. R., Deyo, R. A., & Disterhoft, J. F. (1990). Hippocampectomy disrupts trace eye-blink conditioning in rabbits. Behavioral Neuroscience, 104(2), 243–252. 10.1037//0735-7044.104.2.243

Niediek, J., Boström, J., Elger, C. E., & Mormann, F. (2016). Reliable analysis of single-unit recordings from the human brain under noisy conditions: tracking neurons over hours. PLOS ONE, 11(12), e0166598. 10.1371/journal.pone.0166598

Oostenveld, R., Fries, P., Maris, E., & Schoffelen, J.-M. (2011). FieldTrip: Open source software for advanced analysis of MEG, EEG, and invasive electrophysiological data. Computational Intelligence and Neuroscience, 2011, 156869. 10.1155/2011/156869

Puhger, K., Crestani, A. P., Diniz, C. R. A. F., & Wiltgen, B. J. (2024). The hippocampus contributes to retroactive stimulus associations during trace fear conditioning. iScience, 27(3). 10.1016/j.isci.2024.109035

Qin, S., Piekema, C., Petersson, K. M., Han, B., Luo, J., & Fernández, G. (2007). Probing the transformation of discontinuous associations into episodic memory: An event-related fMRI study. Neuroimage, 38(1), 212–222.

Quiroga, R. Q., Reddy, L., Kreiman, G., Koch, C., & Fried, I. (2005). Invariant visual representation by single neurons in the human brain. Nature, 435(7045), 1102–1107. 10.1038/nature03687

Sederberg, P. B., Schulze-Bonhage, A., Madsen, J. R., Bromfield, E. B., McCarthy, D. C., Brandt, A., Tully, M. S., & Kahana, M. J. (2007). Hippocampal and neocortical gamma oscillations predict memory formation in humans. Cerebral Cortex, 17(5), 1190–1196. 10.1093/cercor/bhl030

Solomon, P. R., Vander Schaaf, E. R., Thompson, R. F., & Weisz, D. J. (1986). Hippocampus and trace conditioning of the rabbit’s classically conditioned nictitating membrane response. Behavioral Neuroscience, 100(5), 729–744. 10.1037//0735-7044.100.5.729

Staresina, B. P., & Davachi, L. (2009). Mind the gap: Binding experiences across space and time in the human hippocampus. Neuron, 63(2), 267–276. 10.1016/j.neuron.2009.06.024

Staresina, B. P., Michelmann, S., Bonnefond, M., Jensen, O., Axmacher, N., & Fell, J. (2016). Hippocampal pattern completion is linked to gamma power increases and alpha power decreases during recollection. eLife, 5, e17397. 10.7554/eLife.17397

Staresina, B. P., Reber, T. P., Niediek, J., Boström, J., Elger, C. E., & Mormann, F. (2019). Recollection in the human hippocampal-entorhinal cell circuitry. Nature Communications, 10(1), 1503.

Staresina, B. P., & Wimber, M. (2019). A neural chronometry of memory recall. Trends in Cognitive Sciences, 23(12), 1071–1085.

Stokes, M. G. (2015). ‘Activity-silent’ working memory in prefrontal cortex: A dynamic coding framework. Trends in Cognitive Sciences, 19(7), 394–405. 10.1016/j.tics.2015.05.004

Teyler, T. J., & DiScenna, P. (1986). The hippocampal memory indexing theory. Behavioral Neuroscience, 100(2), 147.

Treder, M. S., Charest, I., Michelmann, S., Martín-Buro, M. C., Roux, F., Carceller-Benito, F., Ugalde-Canitrot, A., Rollings, D. T., Sawlani, V., Chelvarajah, R., Wimber, M., Hanslmayr, S., & Staresina, B. P. (2021). The hippocampus as the switchboard between perception and memory. Proceedings of the National Academy of Sciences of the United States of America, 118(50), e2114171118. 10.1073/pnas.2114171118

Treves, A., & Rolls, E. T. (1994). Computational analysis of the role of the hippocampus in memory. Hippocampus, 4(3), 374–391. 10.1002/hipo.450040319

Tulving, E. (1983). Elements of Episodic Memory. Oxford University Press.

Tulving, E. (2002). Episodic memory: From mind to brain. Annual Review of Psychology, 53, 1–25. 10.1146/annurev.psych.53.100901.135114

Wallenstein, G. V., Eichenbaum, H., & Hasselmo, M. E. (1998). The hippocampus as an associator of discontiguous events. Trends in Neurosciences, 21(8), 317–323. 10.1016/s0166-2236(97)01220-4

Whittington, J. C. R., Muller, T. H., Mark, S., Chen, G., Barry, C., Burgess, N., & Behrens, T. E. J. (2020). The tolman-eichenbaum machine: Unifying space and relational memory through generalization in the hippocampal formation. Cell, 183(5), 1249–1263.e23. 10.1016/j.cell.2020.10.024

Wolff, M. J., Jochim, J., Akyürek, E. G., & Stokes, M. G. (2017). Dynamic hidden states underlying working-memory-guided behavior. Nature Neuroscience, 20(6), 864–871. 10.1038/nn.4546

Zhu, Y., Zeng, Y., Ren, J., Zhang, L., Chen, C., Fernandez, G., & Qin, S. (2022). Emotional learning retroactively promotes memory integration through rapid neural reactivation and reorganization. Elife, 11, e60190.

